# The Impact of Glycosylation on the Conformational Ensembles of *β*-, *δ*-, and *γ* Sarcoglycans

**DOI:** 10.64898/2026.01.13.699269

**Authors:** Elham Fazelpour, Gabriel A. Cook, Martin McCullagh

## Abstract

Glycosylation is a pivotal post-translational modification that influences protein folding, stability, and interactions, with direct implications for muscular dystrophy pathogenesis and emerging gene therapies. Sarcoglycans (SGs), *β*-, *δ*-, *γ*-, and *α*- subunits of the dystrophin–glycoprotein complex (DGC), contain essential N-linked glycosylation sites, and mutations that disrupt glycan attachment, destabilize the complex and cause limb-girdle muscular dystrophy. Yet, the structural consequences of SG glycosylation remain poorly defined due to the absence of experimental sarcoglycan complex structures. Here, we use homology modeling, AlphaFold predictions, and allatom molecular dynamics simulations to probe how N-linked glycans reshape the conformational ensembles of *β*-, *δ*-, and *γ*-SG monomers and the *β*–*δ*–*γ* heterotrimer core. We find that glycosylation increases flexibility and conformational heterogeneity in isolated monomers but reinforces a compact, stabilized architecture in the heterotrimer. Contact map and clustering analyses show that glycans redistribute local residue interactions while preserving global trimer organization, suggesting a context-dependent role in destabilizing monomers yet reinforcing complex stability. These findings provide the first atomistic insight into how glycosylation primes sarcoglycans for assembly and may explain why mutations at glycosylation sites disrupt complex integrity and drive muscular dystrophy phenotypes.

## 1 Introduction

Sarcoglycans (SGs) are single-pass transmembrane glycoproteins that form an essential sub-complex of the dystrophin–glycoprotein complex (DGC), where they maintain muscle cell integrity and transmit mechanical signals during contraction.^1,2^ Mutations in SG genes destabilize this sub-complex and cause autosomal recessive limb-girdle muscular dystrophies (LGMD), with the loss of any one subunit often leading to degradation of the entire complex.^3^ N-linked glycosylation is introduced during folding in the endoplasmic reticulum and further processed in the Golgi, and is critical for SG stability, trafficking, and membrane localization. Clinical observations underscore this importance: for example, mutations in *α*-SG (e.g., R77C) allow partial retention of the *β*–*δ*–*γ* core at the membrane, whereas loss of *β*- or *δ*-SG abolishes complex formation.^4,5^ These findings highlight the central role of the *β*–*δ*–*γ* core in complex stability and provide a rationale for focusing on this trimer in structural and mechanistic studies.

The structural organization of the *β*–*δ*–*γ* trimer reveals how these subunits form a stable platform for sarcoglycan assembly. Together, these type II transmembrane proteins adopt a boomerang-shaped architecture with three distinct regions: the “grip,” “arm,” and “head.” The N-terminal transmembrane (TM) domains form the grip, a twisted three-helix bundle stabilized primarily by hydrophobic interactions. Each subunit contains a conserved asparagine residue that mediates specific interhelical contacts (N79, N48, and N50 in *β*-, *δ*-, and *γ*-SG, respectively; Figure 1a). Beyond the membrane, extracellular *β*-strands from all three subunits co-fold into a rigid *β*-helix arranged in a staggered, spiral-like fashion, producing a tightly interlocked triangular architecture. The C-terminal loops of each monomer, containing conserved disulfide bonds, further stabilize both the individual subunits and the trimer by forming backbone contacts across subunits. Altogether, these cooperative interactions maintain the cohesion of the *β*–*δ*–*γ* core, providing a scaffold for subsequent recruitment of *α*-SG.^6^

**Figure 1:**
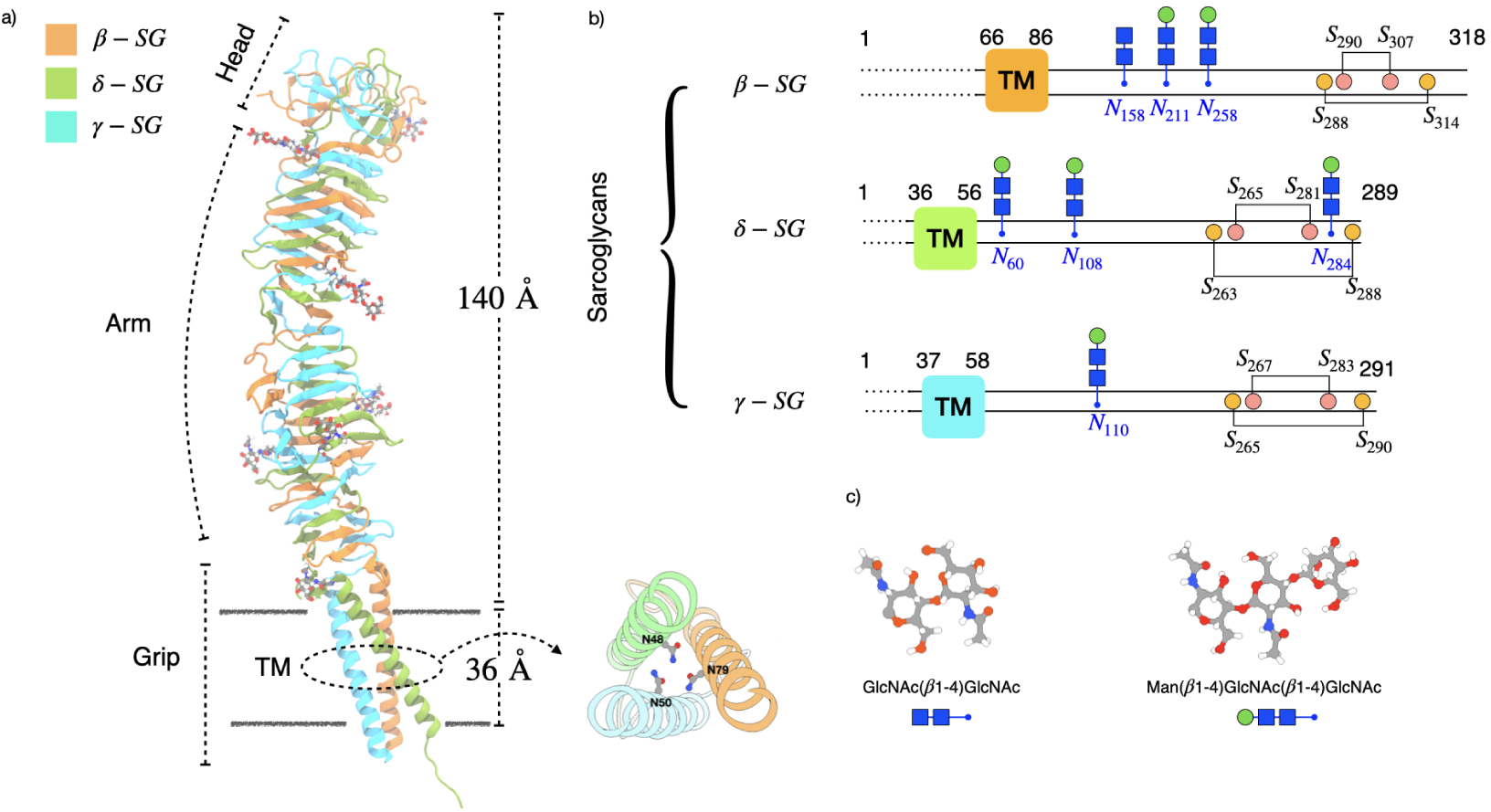
Triple *β*-helix structure of heterotrimer Sarcoglycans. a) A boomerang-like *β*- helix formed by *β*-SG–*δ*-SG–*γ*-SG. Left, the section names of the boomerang-like *β*-helix are labeled. “Head” and “Arm” section construct the extracellular region while “Grip” contains the Transmembrane (TM) and Intracellular regions. Bottom right highlights a detailed extracellular-facing view of the transmembrane domain. b) Domain arrangement of heterotrimer SG complex. Residue numbers at domain boundaries are indicated. Modeled *N* -glycan sites and the sugar structures are shown in blue. Disulfide bond pairs are shown in yellow and pink. Unresolved or missing residues are indicated by dashed lines for SGs. c) The sugar structures along with their glycan code used at *N* -glycan sites.

**Figure 2:**
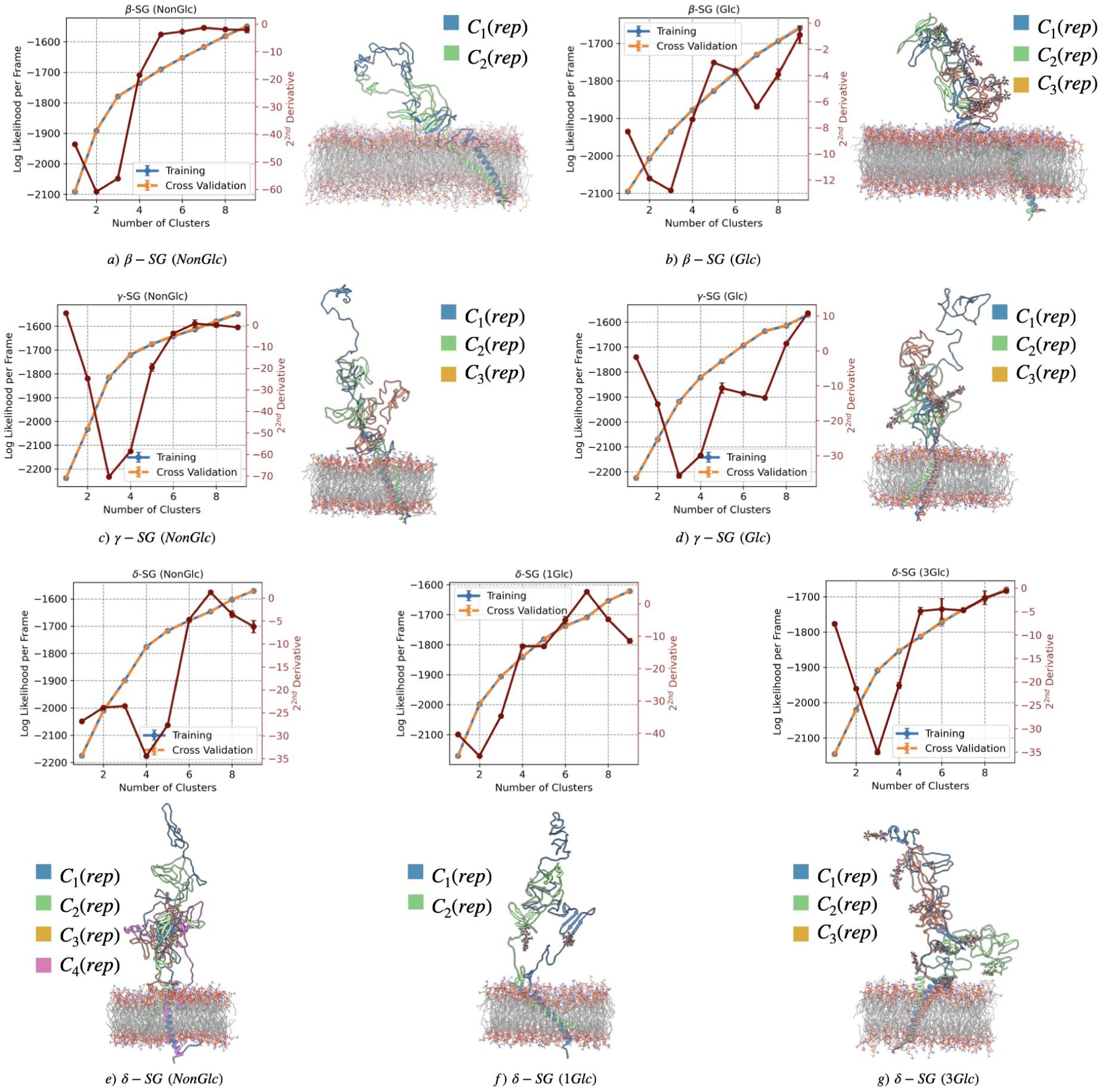
Identification of unique protein conformational clusters from amalgamated trajectories of monomeric *β*-SG, *γ*-SG, and *δ*-SG in glycosylated (Glc) and nonglycosylated (NonGlc) states (a–g). Each plot depicts the log likelihood per frame as a function of the number of clusters for the uniform shape-GMM. Two curves are shown in each plot: the training set (blue) and the cross-validation (CV) set (orange). Error bars represent the standard deviation obtained from sampling 10 different training sets. For each system, the representative protein structure of a cluster component is also shown and color coded accordingly, with the number of clusters ranging from a minimum of 2 to a maximum of 4. The protein structures are superimposed to better represent the conformational differences.

Despite its central role, the contribution of glycosylation to sarcoglycan folding, stability, and assembly remains poorly defined. This knowledge gap is particularly concerning in light of recent safety issues in muscular dystrophy gene therapies, such as the FDA’s suspension of Sarepta Therapeutics’ Elevidys trials following patient deaths from acute liver failure.^7^ Moreover, no experimental structure of the human SG complex currently exists, and available models are limited to isolated domains or computational predictions.^8,9^ How glycosylation modulates the conformational landscape and inter-subunit interactions of *β*-, *δ*-, and *γ*-SG remains largely unknown, representing a critical barrier to mechanistic understanding and therapeutic design.

Glycosylation is well established to influence protein function by altering the conformational ensemble. In sarcoglycans, mutation-induced changes such as the R71T substitution in *δ*-SG introduce new N-glycan sites and shift molecular weight, consistent with structural perturbations, while enzymatic removal of glycans from *γ*- and *δ*-SG alters apparent size and stability.^10^ These findings suggest that glycans can remodel local folding preferences and inter-subunit contacts, thereby influencing complex assembly and function. The structural and functional consequences of N-linked glycosylation in sarcoglycans remain incompletely understood. Experimental studies suggest that glycans can alter folding stability, dynamics, and inter-subunit interactions; however, the lack of high-resolution structural data for the human sarcoglycan (SG) complex has limited mechanistic insight into these effects. To address this gap, we combine homology modeling and AlphaFold-based structural predictions with extensive all-atom molecular dynamics simulations of *β*-, *δ*-, and *γ*-SG monomers and their heterotrimeric core complex. We hypothesize that glycosylation modulates sarcoglycan conformational ensembles by redistributing metastable states and buffering local perturbations upon complex formation. In isolated subunits, glycans increase conformational heterogeneity and sensitivity to perturbation. In contrast, within the assembled *β*–*δ*–*γ* trimer, glycosylation stabilizes dominant conformational states and buffers structural fluctuations, thereby pre-organizing the core complex for robust assembly and function.

## 2 Computational Methods

### 2.1 Starting Structures

Currently, no experimental structural data are available for the human sarcoglycan complex. To model this structure, homologs from *Mus musculus* (PDB ID: 8YT8) and *Oryctolagus cuniculus* (PDB ID: 9C3C), which share high sequence identities with the human counterpart (91.66% and 94.54%, respectively),^8,9^ were used as templates. Guided by these homologs, the amino acid sequences of *β*-, *δ*-, and *γ*-sarcoglycans were extracted from UniProt^11^ (see Table S1) and modeled as both monomers and a heterotrimer using AlphaFold^12,13^ and homology modeling via the SWISS-MODEL server.^14,15^ To mimic the *in vitro* environment, all structures, whether as a monomeric or heterotrimeric system, were integrated into a DMPC bilayer membrane (1,2-dimyristoyl-sn-glycero-3-phosphocholine).^16^ Simulation details differed between monomers and the heterotrimer and are discussed separately.

### 2.2 SG monomeric Subunit simulation preparation

Glycosylated and non-glycosylated systems of *β*-, *δ*-, and *γ*-sarcoglycan were modeled using CHARMM36m parameters.^17,18^ Simulations were prepared with the CHARMM-GUI server,^19,20^ and each subunit was oriented according to the Orientations of Proteins in Membranes (OPM) database,^21^ with standard N- and C-terminal patches applied. Each SG subunit was embedded in a separate DMPC bilayer, solvated with TIP3P water molecules to provide a minimum 15 Å buffer above and below the membrane, and neutralized with Na^+^ and Cl*^−^* ions to achieve a 0.1 M salt concentration. For glycosylated systems, CHARMM-GUI protocols were used to apply glycans at reported sites during the PDB modification step. Among the three SG subunits, *δ*-SG is unique in having two possible glycosylation sets: one experimentally reported site (N108, 1Glc) and three predicted sites (N60, N108, and N284, 3Glc). Details of the starting structures and simulation protocols are provided in Tables S1 and S2 of the supporting document.

### 2.3 SG heterotrimer simulation preparation

The sarcoglycan heterotrimer complex, consisting of *β*-, *δ*-, and *γ*-SG, was also modeled using the same protocol. For the heterotrimer, two glycosylated systems were generated corresponding to the two possible glycosylation sets of *δ*-SG. In the first, *δ*-SG was glycosylated only at N108, giving a total of five glycosylation sites in the complex (5Glc). In the second, *δ*-SG was glycosylated at all three predicted sites, giving a total of seven glycosylation sites (7Glc; see Table S3).

### 2.4 Molecular Dynamics Simulation Protocol

All-atom, explicit solvent molecular dynamics (MD) simulations for glycosylated and non-glycosylated SG monomeric subunits (*β*-SG, *δ*-SG and *γ*-SG) and SG heterotrimer complex were executed using the AMBER18 package.^17^ The cutoff distance for non-bonded interactions was set to 12 Å, after which Coulombic interactions were treated with the particle mesh Ewald method.^22^ The effects of long-range van der Waals interactions were estimated using a dispersion correction model. Periodic boundary conditions were employed. Using the equilibration and production inputs generated by CHARMM-GUI, we first ran the established seven-step minimization/equilibration process.^23,24^ Then, 20 independent 100 ns production runs were performed using the GPU-accelerated CUDA^25^ version of AMBER18, pmemd.^26^

The initial systems were minimized for 5000 steps. Positional restraints were applied to the protein, sugars, ligands, and lipid head groups with a force constant of 10.0 kcal mol*^−^*^1^Å*^−^*^2^, while dihedral restraints were applied to sugars and lipids. Positional restraints were defined for specific residues and groups, ensuring efficient system minimization while maintaining the structural integrity of key protein and membrane regions. Then, equilibration was done using a multi-step process to gradually reduce positional restraints and stabilize the system before the production run. The initial equilibration step was over 1 ns NVT simulation, which used a high positional restraint of 250.0 kcal mol*^−^*^1^Å*^−^*^2^ to stabilize the system while allowing the solvent to equilibrate around the protein and membrane. Over the next five equilibration steps, for a total of 8 ns, the positional restraint force constants were gradually reduced in the following order: 250.0, 100.0, 50.0, 50.0, and 25.0 kcal mol*^−^*^1^Å*^−^*^2^. This gradual reduction helps to maintain the structural integrity of the protein and other molecules while allowing the solvent and ions to equilibrate around them.

For the production simulations, we used the standard input file for NPT simulations generated by CHARMM-GUI, with temperature control using the Langevin thermostat^27^ with a friction coefficient of 1.0 ps*^−^*^1^ and semi-isotropic pressure control using the Berendsen barostat^28^ with a relaxation time of 1.0 ps. The target temperature and pressure were set to 298.15 K and 1.0 bar, respectively. Production runs of both glycosylated and nonglycosylated SG simulations (SG complex and subunits) were performed in triplicate for 1 *µ*s each, yielding a total of 3 *µ*s for each system.

### 2.5 Analyses

Local and global attributes of the conformational ensembles were extracted from the MD simulations. Standard analyses such as principal component analysis, RMSD, RMSF, and DSSP are described in Supporting Information (Section S1).

#### 2.5.1 Conformational clustering

In this study, we employed size-and-shape space Gaussian Mixture Model (Shape-GMM)^29^ to identify structural states (clusters) of SG proteins (individual subunits and complex) based on particle positions. The model fits multivariate Gaussian distributions to the data and estimates optimal parameters for each cluster. To determine the appropriate number of clusters, we used the elbow heuristic method alongside cross-validation (CV). The elbow heuristic method identifies the point at which adding more clusters yields diminishing improvement in log-likelihood, typically marked by a minimum in its second derivative. Cross-validation was used to prevent overfitting, with five training sets generated to estimate sampling error. Frames were assigned to clusters by minimizing the Mahalanobis distance following uniform alignment. All Shape-GMM analyses used a uniform product model for the covariance matrices.

#### 2.5.2 NMR Chemical Shift Calculations

In this work, only the *γ*-SG structure experimental NMR data was available to assess the quality of the predicted structure.^16^ Given this, SHIFTX2^30^was used to compute the backbone and side chain ^1^H and ^15^N chemical shifts for *γ*-SG using the last 50ns of the trajectory from molecular dynamics simulations across all three replicas (Figure S1).

## 3 Results and Discussion

N-linked glycosylation modulates protein folding and stability through a variety of mechanisms.^31–33^ In the case of large glycoproteins like sarcoglycans, glycans typically influence local conformational preferences and dampen fluctuations.^34^ In this study, we aim to investigate the impact of glycosylation on the conformational ensemble of SG complex. Since glycosylation-sites are located on the ECD region and most of the interactions happen at that region, in most of the performed analyses we only consider the residues in extracellular region. To accomplish this, we analyze simulations of glycosylated (Glc) and non-glycosylated (NonGlc) of *β*-, *δ*- and *γ*-SG as an isolated structure (or monomer) and heterotrimer complex. We subsequently cluster conformational ensembles using shape-GMM. The changes in the number of conformational clusters observed in monomeric sarcoglycan subunits may influence how these subunits participate in sarcoglycan complex assembly.

### 3.1 Glycosylation impact on Monomers

#### Model Corroboration

To examine the influence of glycosylation on the conformational ensemble of each monomer, we initially employed SHIFTX2 NMR prediction algorithm to validate the predicted *γ*-SG structure (Figure S1); this analysis was limited to *γ*-SG, as experimental data were available only for this subunit.^16^ Subsequently, shape-GMM clustering analysis followed by PCA analysis were performed on both glycosylated and non-glycosylated monomers to investigate the impact of glycosylation on the diversity of conformational states. Finally, Root-mean-square fluctuations (RMSFs), secondary structure (DSSP) and residue contact map analyses were conducted to identify residue-level changes induced by glycosylation.

The comparison of experimental NMR data of non-glycosylated *γ*-SG with predicted chemical shifts reveals a generally acceptable alignment across the protein’s structure. The data in Figure S1, segmented into Intra Cellular Domain (ICD), Transmembrane Domain (TMD), and Extracellular Domain (ECD) (see Table S1), alongside a full structure overview, shows that predicted shifts cluster around 115-120 ppm (N Chemical Shift) and 6-8 ppm (H Chemical Shift), with the highest density (yellow-green) matching the concentration of experimental points (purple dots). Notably, the TMD exhibits the best agreement, with a distinct predicted cluster closely overlapping the experimental data, which is not surprising given that the helical TMD region is embedded and restrained within the DMPC bilayer membrane. The ICD, with fewer experimental points, and the broader ECD, with a wider distribution, also show reasonable consistency, though some experimental points fall outside high-density predicted areas, potentially indicating structural flexibility, post-translational modifications, or limitations in the SHIFTX2 model. Overall, SHIFTX2 provides a reliable approximation of the NMR chemical shifts for *γ*-SG, with the full structure analysis reinforcing this trend. The observed discrepancies, particularly in less densely sampled or more variable regions, highlight areas for further investigation, potentially requiring additional experimental data or refined predictive models.

#### The Ensemble view

Glycosylation reshapes the conformational ensemble by altering both the number and nature of accessible structural states. To quantify these effects, amalgamated trajectories from three replicas of each system were analyzed using conformational clustering, followed by PCA on aligned coordinates to capture the dominant modes of structural variation.

For *β*-SG, clustering reveals pronounced differences between non-glycosylated and glycosylated systems. The NonGlc form populates two major clusters, C_1_ (76.05%) and C_2_ (23.94%), separated by a large RMSD of 11.06Å indicating distinct conformational states. In contrast, the Glc form samples three major clusters with substantial RMSD separation between representatives (Table 1). PCA further highlights these differences: the NonGlc system exhibits two well-separated, high-density regions broadly distributed along PC1 and PC2, reflecting high flexibility and multiple metastable states (Figure 3.a). The Glc system instead displays a more compact and continuous density forming a central basin, with smoother transitions and a narrower ensemble of stable conformations (Figure 3.b).

**Figure 3:**
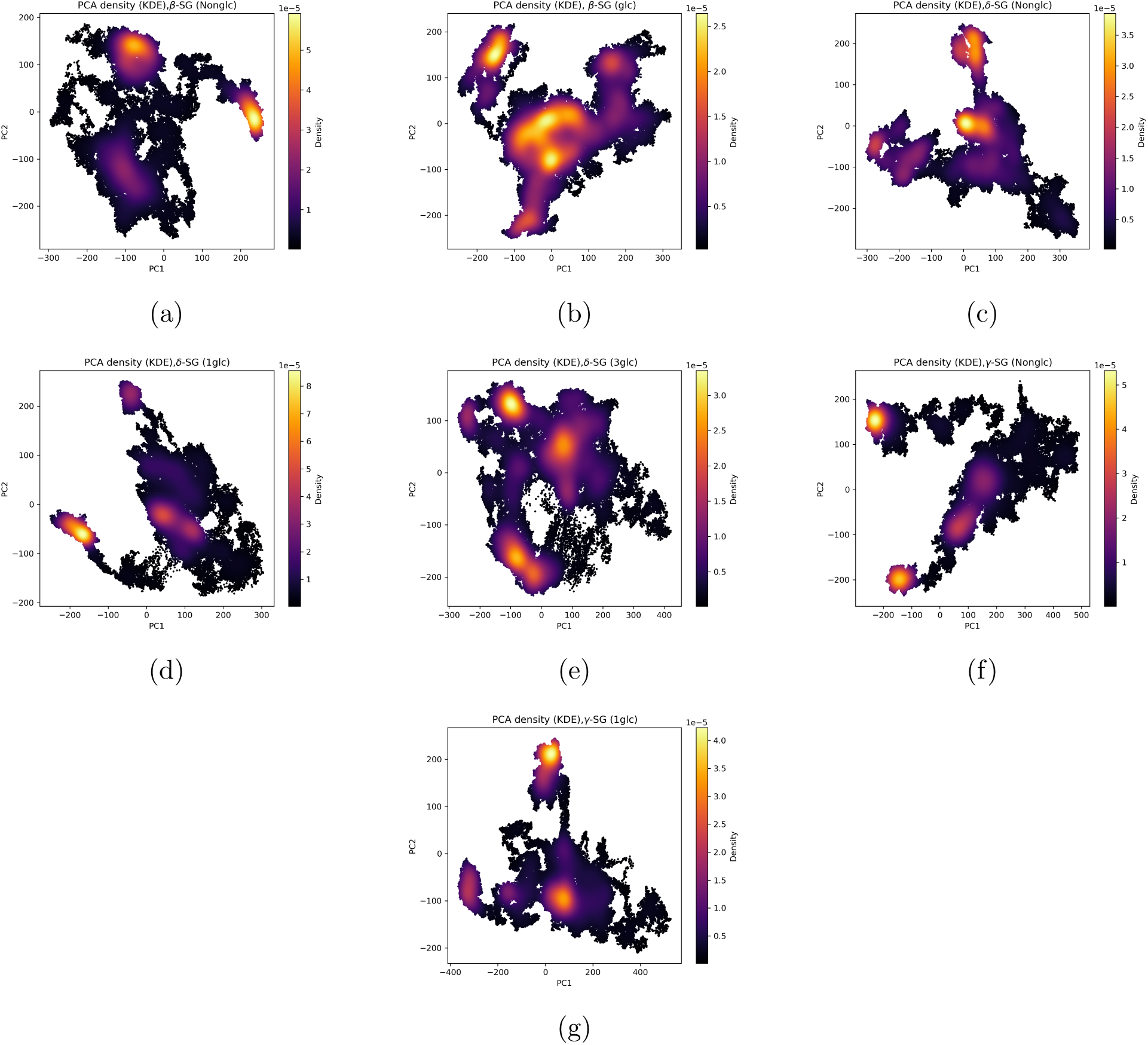
Two-dimensional principal component analysis (PCA) kernel density estimates (KDEs) illustrating the conformational sampling of non-glycosylated and glycosylated *β*-, *δ*-, and *γ*- sarcoglycan monomers. Each panel represents the projection of atomic coordinates onto the first two principal components obtained from aligned MD trajectories. The color intensity reflects the probability density of sampled conformations.

**Table 1:** Relative population of clusters in Monomeric sarcoglycan system calculated using Shape-GMM.

| Name | Population |  |  |  |
| --- | --- | --- | --- | --- |
|  | C<sub>1</sub>(%) | C<sub>2</sub>(%) | C<sub>3</sub>(%) | C<sub>4</sub>(%) |
| β-SG | 76.05 | 23.94 | | |
| β-SG (Glc) | 54.22 | 28.34 | 17.42 | |
| δ-SG | 36.47 | 24.18 | 23.35 | 16.00 |
| δ-SG (1Glc) | 72.84 | 27.15 | | |
| δ-SG (3Glc) | 53.43 | 24.48 | 22.07 | |
| γ-SG | 65.11 | 24.06 | 19.81 | |
| γ-SG (Glc) | 57.91 | 28.57 | 13.51 | |

**Table 2:** RMSD(*Å*) between the representative of each cluster in monomeric sarcoglycan systems.

| Name | RMSD($\text{\AA}$) | | | | | |
| --- | --- | --- | --- | --- | --- | --- |
|  | C<sub>1</sub>-C<sub>2</sub> | C<sub>1</sub>-C<sub>3</sub> | C<sub>1</sub>-C<sub>4</sub> | C<sub>2</sub>-C<sub>3</sub> | C<sub>2</sub>-C<sub>4</sub> | C<sub>3</sub>-C<sub>4</sub> |
| β-SG | 11.063 | | | | | |
| β-SG (Glc) | 8.4977 | 9.309 | | 12.2494 | | |
| δ-SG | 11.3263 | 9.7025 | 8.5552 | 12.5851 | 11.2813 | 10.0565 |
| δ-SG (1Glc) | 9.3704 | | | | | |
| δ-SG (3Glc) | 9.4443 | 8.9245 | | 11.1242 | | |
| γ-SG | 10.9289 | 14.5942 | | 16.3653 | | |
| γ-SG (Glc) | 14.3503 | 12.9259 | | 13.5235 | | |

For *δ*-SG, glycosylation progressively restricts conformational diversity. The NonGlc system exhibits four distinct clusters and a broad PCA distribution with multiple high-density regions (Figure 3.c), indicative of a diverse conformational ensemble. Addition of one glycosylation site (1Glc) reduces the system to two dominant clusters and produces a more concentrated density distribution (Figure 3.d). Further glycosylation (3Glc) results in three clusters and an even more localized PCA density (Figure 3.e). These trends indicate increasing rigidity and stabilization with increasing glycosylation.

For *γ*-SG, both Glc and NonGlc systems exhibit three major clusters with large intercluster RMSDs (Table 1), yet their conformational landscapes differ markedly. PCA reveals multimodal distributions characteristic of heterogeneous sampling; however, the glycosylated form displays a compact, high-density basin centered near the origin, consistent with N-linked glycan–mediated stabilization of the extracellular domain (Figure 3.g). In contrast, the NonGlc system exhibits a broader, more diffuse distribution with extended sampling along PC1, indicating increased flexibility and reduced conformational stability (Figure 3.f). This behavior is consistent with experimental evidence linking incomplete glycosylation to sarcoglycan destabilization and LGMD2C pathogenesis.^35^ Overall, glycosylation compacts the conformational space of *γ*-SG, stabilizing its native fold and supporting the structural integrity of the sarcoglycan complex.

#### Glycosylation impact on secondary structure

DSSP analysis was used to assess glycosylation-induced local secondary structure changes. For *β*-SG, the overall distributions of *α*-helices, *β*-sheets, and coils are similar in Glc and NonGlc systems, indicating conserved global secondary structure. However, glycosylation modulates local stability, with residues at or near glycosylation sites (GLY259, SER210, VAL155, SER260, THR209) transitioning from coil to more ordered *β*-sheet or *α*-helical states (Figure 4a). In *δ*-SG, both 1Glc and 3Glc systems exhibit increased persistence of helical and *β*-sheet regions, with the 3Glc system showing the strongest stabilization, consistent with PCA results (Figure 4b and Figure 4c). Glycosylation enhances secondary structure persistence, likely via glycan-induced hydrogen bonding and steric constraints. While residues at glycosylation sites show increased coil content, adjacent regions (e.g., residues 130–162) display enhanced *β*-sheet formation, reflecting a balance between local flexibility and regional stabilization. For *γ*-SG, DSSP reveals pronounced changes in the C-terminal region. Regions near the glycosylation site (N110) and within the C-terminus (residues 239–265) exhibit loss of secondary structure, indicating increased flexibility (Figure 4d).

**Figure 4:**
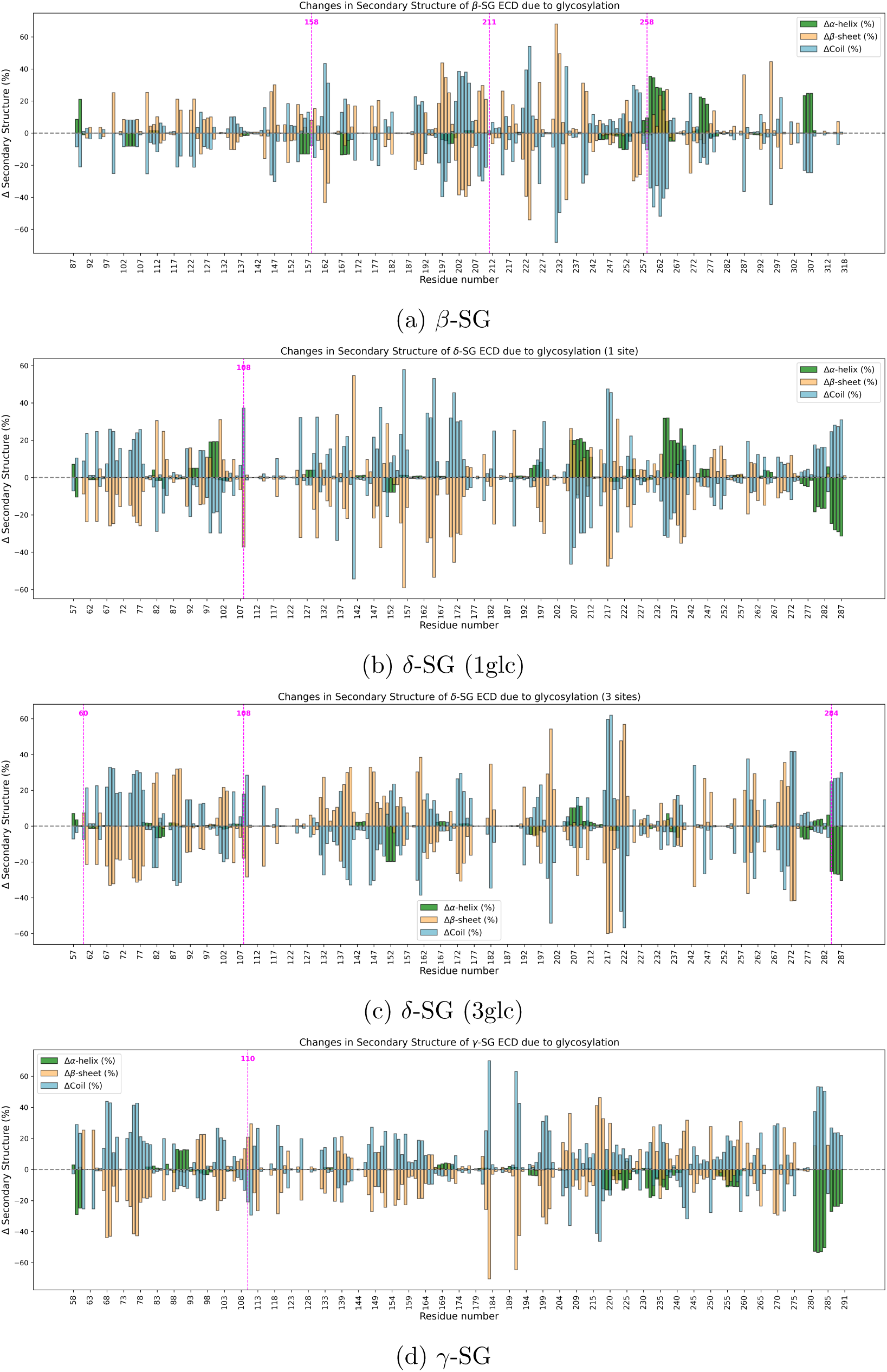
Differences in percentages between glycosylated and non-glycosylated states for *β*, *δ*, and *γ* monomeric SGs. The magenta dashed lines indicate glycosylation sites.

#### Differential localization and conformational effects at residue level

To assess glycosylation-induced changes in intrinsic flexibility, residue-wise RMSF profiles were computed for monomeric *β*-, *δ*-, and *γ*-sarcoglycans (Figure S2). The three isoforms exhibit distinct responses. In *β*-SG, glycosylation causes a modest, global increase in flexibility without evidence of destabilization. In *δ*-SG, glycosylation produces region-specific effects, reducing fluctuations near the termini while increasing mobility in the central region. In contrast, *γ*-SG shows reduced RMSF near the glycosylation site (N110) and the C-terminus, indicating glycan-mediated local stabilization that may support proper folding and complex assembly.

To further resolve residue-level structural effects, contact difference maps were analyzed to identify glycosylation-induced interaction changes beyond global metrics. In monomeric *β*- SG, glycosylation causes widespread contact reorganization, with both gains and losses across multiple regions, indicating substantial local rearrangements (Figure 5a). Similar trends are observed in *δ*-SG, where glycosylation induces extensive contact changes, particularly near glycosylation sites (Figure 5b,c). In *γ*-SG, glycosylation prominently disrupts contacts in the vicinity of the glycan attachment site (Figure 5d).

**Figure 5:**
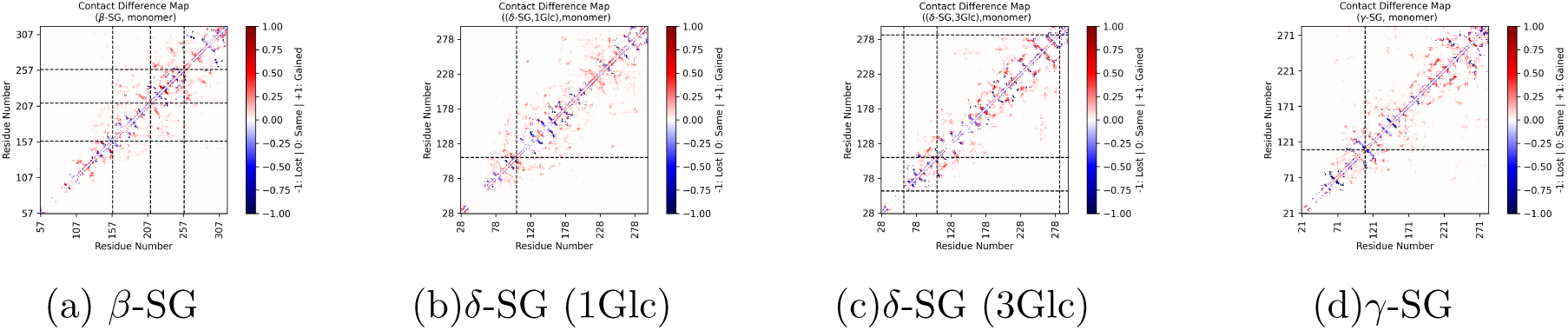
Contact map analysis of *β*-, *δ*- and *γ*-SG as a monomer. Dashed lines indicate glycosylation sites. Green clusters represent new contacts or interactions formed due to glycosylation, while purple clusters indicate contacts or interactions that are lost as a result of glycosylation.

In summary, glycosylation modulates sarcoglycan monomers by reshaping their conformational landscapes, stabilizing dominant structural states, and reorganizing residue-level interactions in an isoform-specific manner. Ensemble analyses show that glycosylation generally compacts the conformational space most strongly in *δ*-SG with increasing glycosylation, while *β*- and *γ*-SG display redistribution and stabilization of distinct states, respectively. Local DSSP, RMSF, and contact-map analyses reveal that glycan attachment alters secondary structure persistence, intrinsic flexibility, and residue interactions, particularly near glycosylation sites, highlighting glycosylation as a key regulator of sarcoglycan structural stability and dynamics.

### 3.2 Glycosylation impacts on heterotrimer SG complex

Given that glycosylation shapes monomer conformations and stability, we next examine its impact on the heterotrimeric *β*–*δ*–*γ* complex. Glycan addition modulates inter-subunit interactions, redistributes structural fluctuations, and stabilizes dominant conformational states, while preserving the overall fold of the complex.

#### Model Corroboration

Given the lack of experimentally resolved structures for the human sarcoglycan complex, the models were constructed using experimentally resolved *Mus musculus* (PDB ID: 8YT8) and *Oryctolagus cuniculus* (PDB ID: 9C3C) homologs as templates, which share high sequence identities of 91.66% and 94.54%, respectively.^8,9^ We evaluated the internal structural stability of the modeled systems using backbone RMSD as a form of model corroboration. The heterotrimeric system displays RMSD convergence after equilibration, with no evidence of progressive structural drift over the production trajectories (Figure S3). The comparable RMSD stability across components of the heterotrimeric complex indicates that the models preserve global fold integrity within the membrane environment, allowing observed differences in conformational heterogeneity and assembly-dependent behavior to be interpreted as intrinsic features of the system rather than consequences of global destabilization.

#### The Ensemble view

The SG complex exhibits a glycosylation-dependent reorganization of its conformational ensemble, transitioning from three clusters in the NonGlc state to two in 5Glc, and back to three in 7Glc (Figure 6a–c; Table 3). In the NonGlc system, multiple high-density regions in the PCA map correspond to the identified clusters, with overlapping densities reflecting a flexible and heterogeneous ensemble (Figure 7.a). Glycosylation at five sites leads to a single dominant high-density basin aligned with two clusters, indicating reduced conformational diversity and merging of previously distinct states (Figure 7.b). In the 7Glc system, the PCA density becomes more focused with a pronounced central peak despite the reappearance of three clusters, suggesting increased rigidity accompanied by subtle conformational sub-states arising from additional steric constraints or intramolecular interactions (Figure 7.c). Overall, glycosylation initially stabilizes the complex by consolidating conformational states (NonGlc → 5Glc), followed by the emergence of nuanced structural diversity at higher glycosylation levels (7Glc). The systematic shift toward lower PC1 and PC2 values with increasing glycosylation indicates transitions between discrete cluster centroids rather than along a continuous conformational pathway.

**Figure 6:**
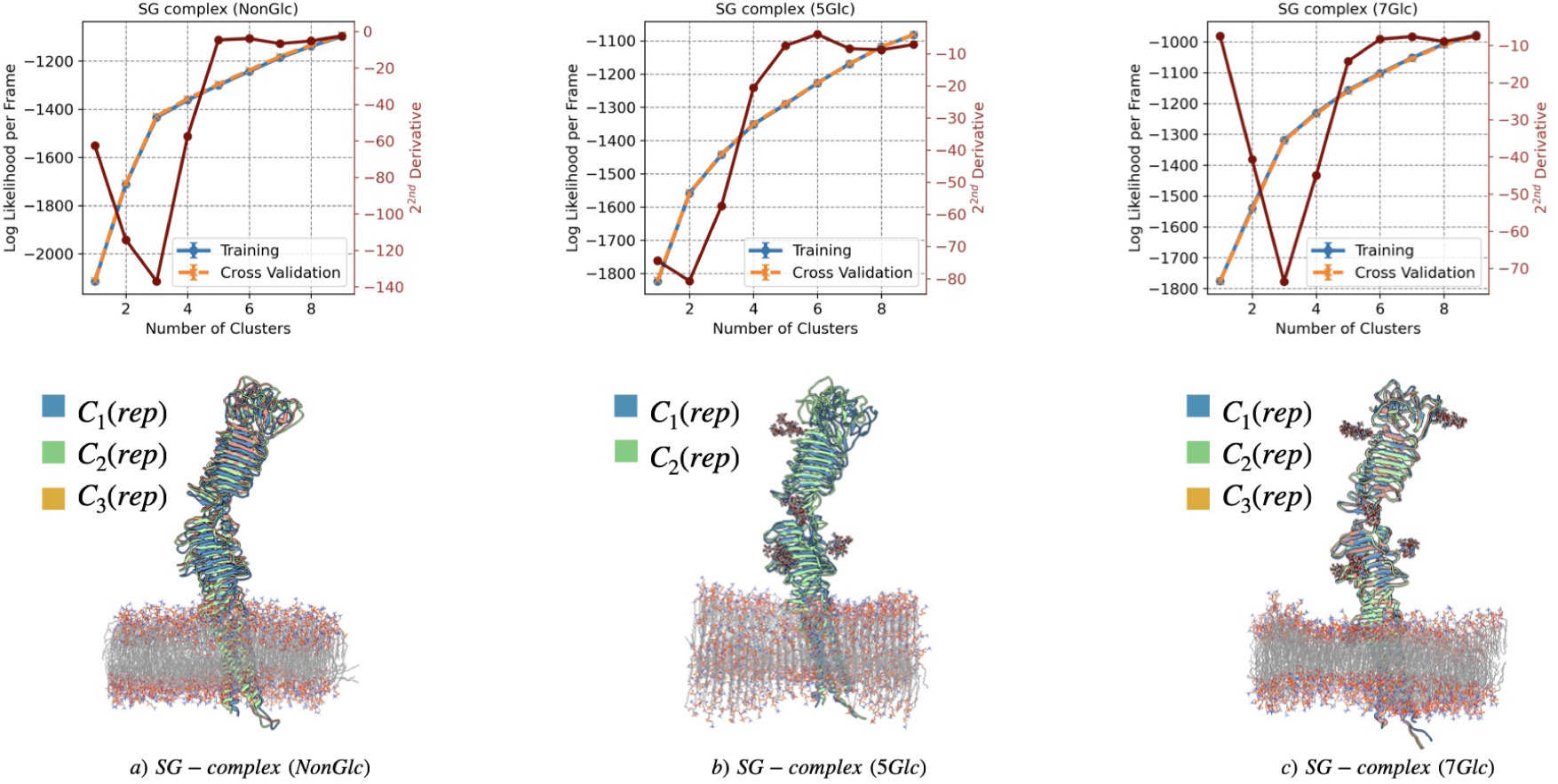
Identification of unique protein conformational clusters from amalgamated trajectories of SG-complex with three subunits. Each plot depicts the log likelihood per frame as a function of the number of clusters for the uniform shape-GMM. Two curves are shown in each plot: the training set (blue) and the cross-validation (CV) set (orange). Error bars represent the standard deviation obtained from sampling 10 different training sets. For each system, the representative protein structure of a cluster component is also shown and color coded accordingly, with the number of clusters ranging from a minimum of 2 to a maximum of 3. The protein structures are superimposed to better represent the conformational differences.

**Figure 7:**
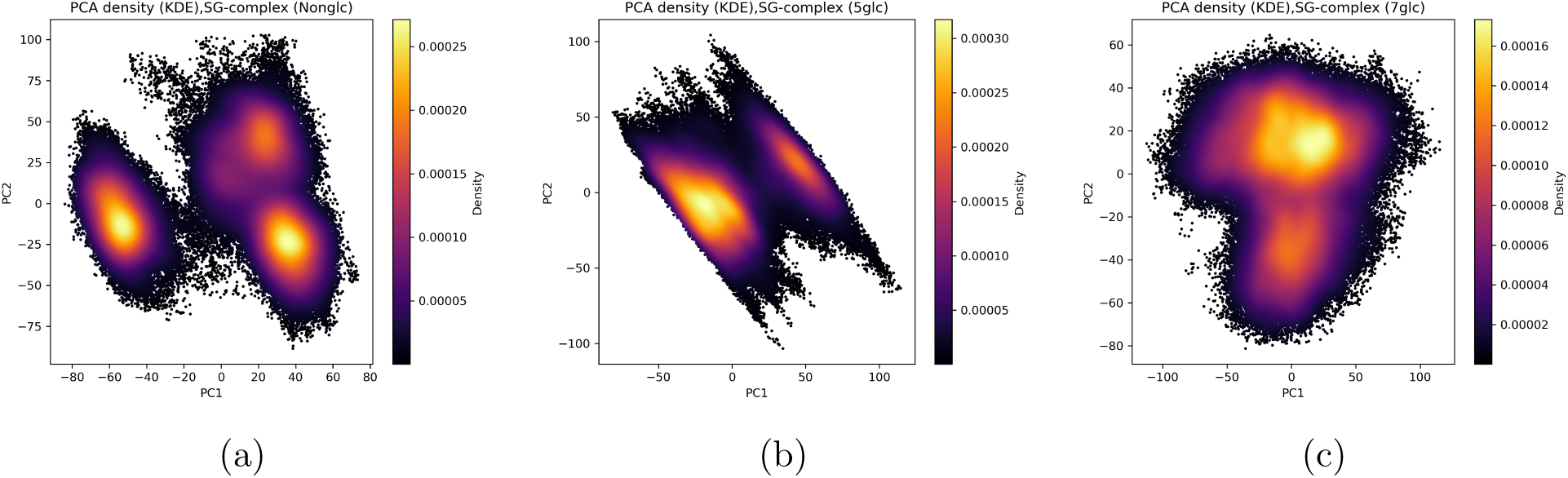
PCA–KDE maps showing conformational sampling of sarcoglycan heterotrimer complex in their non-glycosylated and glycosylated states ( 5 and 7 sites). Color intensity represents conformational density along the first two principal components. Glycosylation modulates the extent and distribution of sampled conformations, indicating isoform-specific dynamic effects.

**Table 3:** Relative population of clusters in heterotrimer sarcoglycan complex systems calculated using Shape-GMM.

| Name | Population |  |  |  |
| --- | --- | --- | --- | --- |
|  | C<sub>1</sub>(%) | C<sub>2</sub>(%) | C<sub>3</sub>(%) | C<sub>4</sub>(%) |
| SG-complex | 34.06 | 33.01 | 32.92 |  |
| SG-complex (5Glc) | 72.18 | 27.81 |  |  |
| SG-complex (7Glc) | 36.58 | 34.59 | 28.81 |  |

**Table 4:** RMSD(Å) between the representative of each cluster in heterotrimer sarcoglycan complex systems.

| Name | RMSD(Å) |  |  |
| --- | --- | --- | --- |
|  | C<sub>1</sub>-C<sub>2</sub> | C<sub>1</sub>-C<sub>3</sub> | C<sub>2</sub>-C<sub>3</sub> |
| SG-complex | 1.3733 | 1.6955 | 1.8987 |
| SG-complex (5Glc) | 1.2725 |  |  |
| SG-complex (7Glc) | 1.2336 | 1.2354 | 1.2937 |

#### Glycosylation impact on Secondary structure

DSSP analysis indicates that secondary structure changes are primarily confined to the C-terminal region, with minimal alterations near glycosylation sites (Figures S3–S5), suggesting that glycosylation does not disrupt the core fold or overall architecture of the SG trimer. While the global structure remains intact, localized glycan-induced effects may still modulate site-specific dynamics or interactions. Overall, the heterotrimeric SG complex is substantially more stable and ordered than the monomeric forms, with glycosylation contributing subtly but meaningfully to trimer stability.

#### Conformational effect at residue level

In the heterotrimeric SG complex, interlocking of *β*-, *δ*-, and *γ*-SG reduces RMSF variability, with further stabilization observed upon glycosylation at five and seven sites (Figure 8), consistent with enhanced inter-subunit interactions.^4,5^ Glycosylation sites (magenta dashed lines) show no pronounced RMSF changes between NonGlc and Glc states in either monomers or the heterotrimer, indicating subtle, localized effects on flexibility. This glycan-mediated stabilization likely influences differential localization by anchoring residues in specific conformations, thereby modulating spatial organization and functional integration within the SG complex.

**Figure 8:**
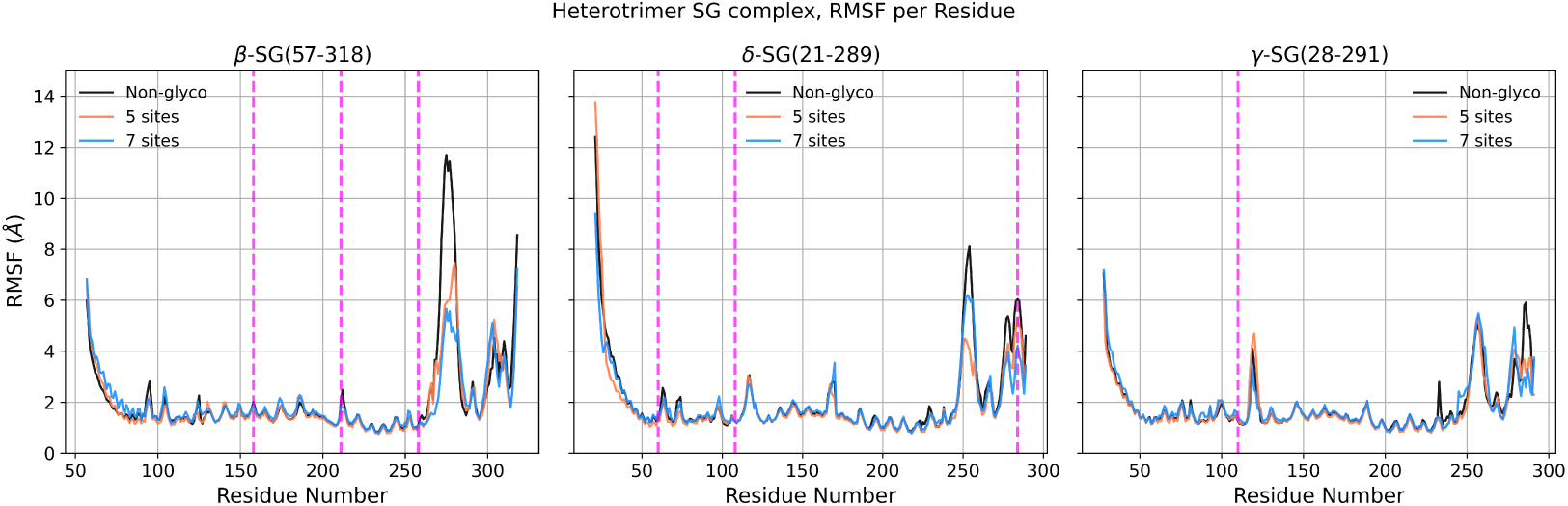
RMSF of SG heterotrimer complex. In each components, C*_α_* of residues was used to calculate the RMSF. The plot on the left belongs to *β*-SG, the one in the middle to *δ*-SG and the one on the right to *γ*-SG.

Compared to monomeric forms, incorporation of *β*-SG into the SG complex with 5Glc results in markedly reduced contact-map variability (Figure 9.a), indicating that inter-subunit interactions buffer glycan-induced perturbations and preserve structural coherence. In contrast, the 7Glc complex shows a pronounced increase in gained off-diagonal contacts (Figure 9.b), suggesting that extensive glycosylation can further reinforce structural cohesion within the complex. These trends highlight a context-dependent role of glycosylation: while glycans reorganize local contacts in isolated monomers, moderate glycosylation stabilizes core interactions in the assembled complex, and higher glycosylation promotes additional inter-residue contacts. These findings are consistent with *β*-SG’s proposed role in initiating sarcoglycan assembly, where glycosylation may enhance local stability while perturbing distant interactions, potentially reflecting compensatory structural mechanisms relevant to LGMD2E.^36,37^ In the SG complex, contact loss is largely confined to regions near glycosylation sites and the C-terminus, likely arising from secondary structure changes or steric effects. By contrast, *δ*- and *γ*-SG exhibit only minor, C-terminal-localized contact changes, reinforcing the stabilizing role of complex assembly in accommodating glycan-induced perturbations (Figure 9.c–f).

**Figure 9:**
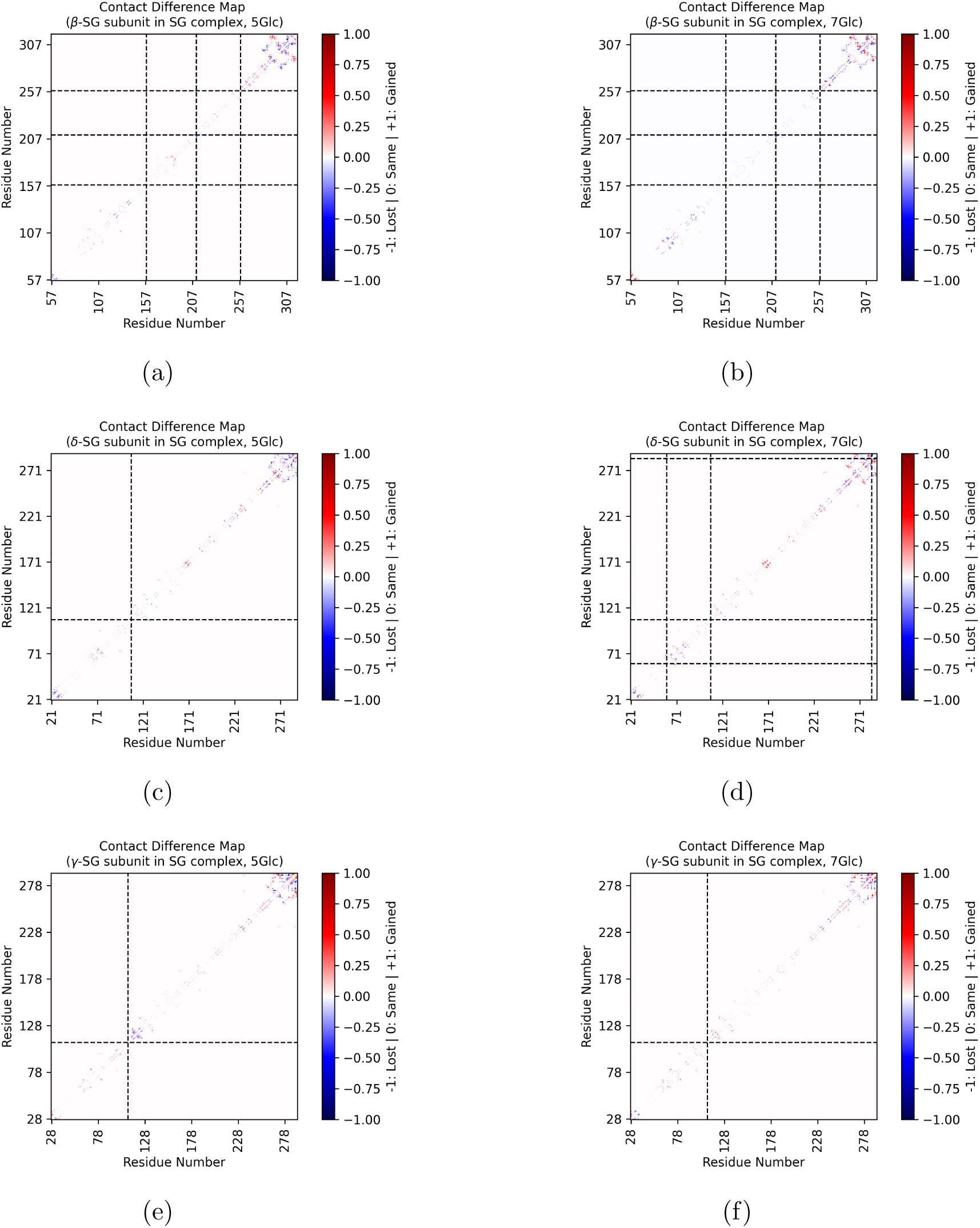
Contact map analysis of *β*-, *δ*- and *γ* subunits of SG. Dashed lines indicate glycosylation sites. Green clusters represent new contacts or interactions formed due to glycosylation, while purple clusters indicate contacts or interactions that are lost as a result of glycosylation.

#### Ensemble changes can have functional implications

N-glycosylation introduces bulky, flexible glycans that modulate protein structure locally and allosterically, often dampening dynamics and influencing regions distal to the modification sites.^38^ Although crystallographic characterization of glycans is challenging, complementary approaches show that glycosylation on flexible loops is particularly sensitive to environmental stress and can strongly affect allosteric communication.^39^ Beyond dynamics, N-glycosylation promotes protein folding and stability by increasing rigidity and protecting against stress, with complex glycans capable of structurally “locking” protein domains into functional orientations.^40,41^ Consistent with these principles, our results indicate that glycosylation modulates conformational heterogeneity in individual sarcoglycan monomers, altering adaptability relevant to complex formation and stress response. Moreover, the SG complex itself likely mediates extracellular-to-intracellular signal transmission,^42–44^ with assembled subunits exhibiting distinct, more regulated flexibility compared to isolated monomers, leading to coordinated local and global structural responses.

## 4 Conclusions

This study demonstrates that N-glycosylation modulates sarcoglycan conformational behavior in a strongly assembly-dependent manner. Isolated SG monomers sample broad and heterogeneous conformational ensembles, reflecting high intrinsic flexibility that renders them sensitive to glycan-induced perturbations. Upon incorporation into the *β*–*δ*–*γ* heterotrimeric complex, conformational variability is markedly reduced, with glycosylation exerting more subtle effects that are buffered by inter-subunit interactions and preserve the overall architecture. Rather than inducing large structural rearrangements, glycosylation shifts the populations of metastable states, tuning local flexibility and residue-level interactions while maintaining the global fold.

These findings suggest that glycosylation acts not as a uniform stabilizer but as a context-dependent regulator that pre-organizes sarcoglycan subunits for assembly, interaction, and functional resilience under stress. By explicitly linking glycan chemistry, conformational dynamics, and assembly state, this work provides a mechanistic framework for understanding how glycosylation defects may selectively destabilize sarcoglycan intermediates and contribute to muscular dystrophy–associated pathologies.

## Acknowledgement

E.F. and M.M acknowledge support from the National Institute for Allergic and Infectious Diseases of the National Institute of Health under award number R01AI166050. Computational resources for this project were provided by the High Performance Computing Center at Oklahoma State University supported in part through the National Science Foundation Grant OAC-1531128.

## Supporting Information

Available

## Notes

### Competing Interest Statement

The authors have declared no competing interest.

