## Supporting Information for "The Impact of Glycosylation on the Conformational Ensembles of *β*-, *δ*-, and *γ* Sarcoglycans"

### The Impact of Glycosylation on the Conformational Ensemble of $\beta$-, $\delta$-, and $\gamma$ Sarcoglycans

#### S1 Analyses

##### S1.1 Secondary Structure

The evolution of secondary structure in glycosylated and non-glycosylated systems of $\beta$-, $\delta$-, and $\gamma$-SG (in separate simulations and as subunit of SG-complex) were analyzed using the DSSP algorithm implemented in the MDTraj Python library.[1] The analysis focused on $\alpha$-helices, $\beta$-sheets, and coil content to assess glycosylation-induced changes. To quantify these changes, we compared the average percentage of each secondary structure element per residue between glycosylated and non-glycosylated systems. These per-residue differences were computed from DSSP output and visualized as bar plots.

##### S1.2 Local Flexibility

Root-mean-square fluctuations (RMSFs) were calculated to assess the average fluctuation of each residue over the trajectory, providing insight into flexibility or mobility changes induced by glycosylation. RMSFs were computed using the *atomicfluct* module in cpptraj, based on the $C_\alpha$ atoms of each residue.[2]

#### S1.3 Contact difference map analysis

All contact map calculations were performed using the MDTraj Python package.[1] $C_\alpha$ atoms corresponding to ECD of the glycosylated and non-glycosylated systems were extracted from each trajectory. All unique $C_\alpha$– $C_\alpha$ residue pairs were enumerated, and pairwise distances were computed for every frame with MDTraj's `compute_distances` function. The mean distance for each pair over the trajectory was calculated and binarized using a 4 Å cutoff, with distances below the cutoff considered contacts. A contact difference map was obtained by subtracting the binary contact matrix of the non-glycosylated system from that of the glycosylated system, yielding values of $-1$ (lost contact), 0 (unchanged), or $+1$ (gained contact). The resulting difference matrix was symmetrized, mapped to UniProt residue numbering of each SG (individually and as a subunit of SG complex), and visualized as a heatmap.

#### S1.4 Principal Component Analysis (PCA) of the Aligned Trajectory

The trajectories were first aligned to remove overall translational and rotational motion prior to performing Principal Component Analysis (PCA). Alignment was performed using the Shape-GMM package, which employs a maximum-likelihood uniform alignment algorithm to superimpose all frames based on the protein's geometry. Using MDAnalysis, $C_\alpha$ atoms of residues were selected to represent the protein backbone, and their Cartesian coordinates were extracted for all frames. The resulting coordinate array (frames $\times$ atoms $\times$ 3) was converted to a PyTorch tensor, centered by removing the center of geometry, and then aligned using `align.maximum_likelihood_uniform_alignment()` to produce a uniformly aligned trajectory. The aligned coordinates were saved as a new trajectory file for downstream analysis. Following alignment, PCA was performed on the aligned coordinates to capture dominant modes of conformational variation. The trajectory was reshaped into a two-dimensional array (frames $\times$ atoms $\times$ 3), and PCA was applied using the scikit-learn implementation.[3] The first two principal components (PC1 and PC2) were used for visualization. Cluster assignments obtained from the Shape-GMM model were mapped onto the PCA projection to examine the relationship between identified conformational clusters and the principal components. Additionally, a kernel density estimate (KDE) was computed using the SciPy `gaussian_kde` function to visualize the density distribution of conformations in PCA space. All analyses and visualizations were performed in Python using MDAnalysis, NumPy, PyTorch, and Matplotlib.[4–7]

### S2 Supporting Tables Figures

Table S1: Summary of model building of the sarcoglycan complex

| subunit | Length(aa) /Uniprot ID | Modeled regions | Domains | Modifications | |
|---|---|---|---|---|---|
| | | | | Glycosylation | Disulfide bonds |
| $\beta$-sarcoglycan | 318 /Q16585 | 57-318 | TM:66-86 ECD:87-318 | N158, N211, N258 | C288-C314 C290-C307 |
| $\delta$-sarcoglycan | 289 /Q92629 | 21-289 | TM:36-56 ECD:57-289 | N60, N108, N284 | C263-C288 C265-C281 |
| $\gamma$-sarcoglycan | 291 /Q13326 | 28-291 | TM:37-58 ECD:59-291 | N110 | C265-C290 C267-C283 |

Table S2: Detailed information about the simulations of both the sarcoglycan subunits and the heterotrimer complex generated using CHARMM-GUI. For the sarcoglycan subunits, the glycosylated and non-glycosylated systems followed the same protocol.

| subunit | Box Size (Å) | # of DMPC | Tilt angle | [NaCl] |
|---|---|---|---|---|
| $\beta$-SG | X:90<br>Y:90<br>Z:222 | 250 | 39° | 0.1M |
| $\delta$-SG | X:90<br>Y:90<br>Z:242 | 250 | 36° | 0.1M |
| $\gamma$-SG | X:90<br>Y:90<br>Z:231 | 250 | 28° | 0.1M |
| heterotrimer complex (non-glycosylated) | X:129<br>Y:129<br>Z:271 | 250 | 1° | 0.1M |
| heterotrimer complex (glycosylated) | X:129<br>Y:129<br>Z:261 | 250 | 1° | 0.1M |

Table S3: Glycan types at each glycosylation site in glycosylated simulations

| subunit | Glycolation site | Glycan type |
|---|---|---|
| $\beta$-SG | N158 | GlcNAc(b1-4)GlcNAc |
| | N211 | Man(b1-4)GlcNAc(b1-4)GlcNAc |
| | N258 | Man(b1-4)GlcNAc(b1-4)GlcNAc |
| $\delta$-SG | N60 | Man(b1-4)GlcNAc(b1-4)GlcNAc |
| | N108 | Man(b1-4)GlcNAc(b1-4)GlcNAc |
| | N284 | Man(b1-4)GlcNAc(b1-4)GlcNAc |
| $\gamma$-SG | N110 | Man(b1-4)GlcNAc(b1-4)GlcNAc |

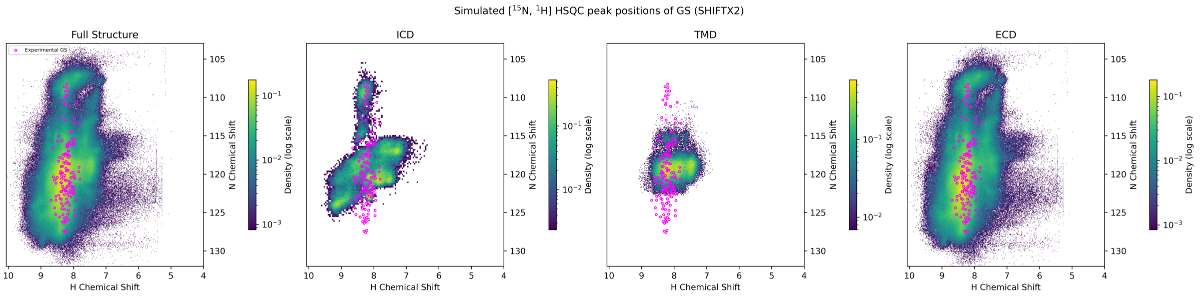


(a) Replica 1

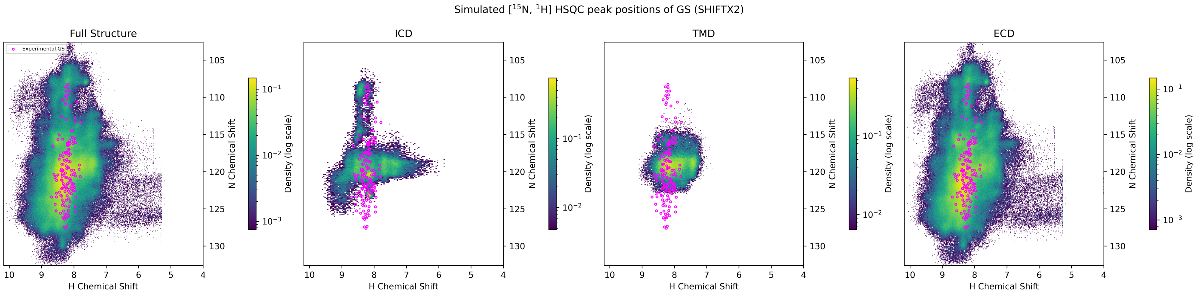


(b) Replica 2

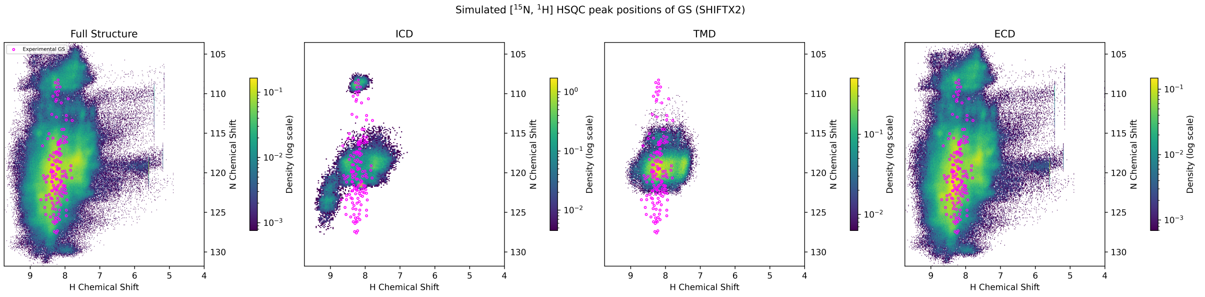


(c) Replica3

Figure S1: Backbone and side chain $^{1}$H and $^{15}$N chemical shifts of $\gamma$-SG using SHIFTX2 and comparison with experimental NMR data (purple dots)[8]

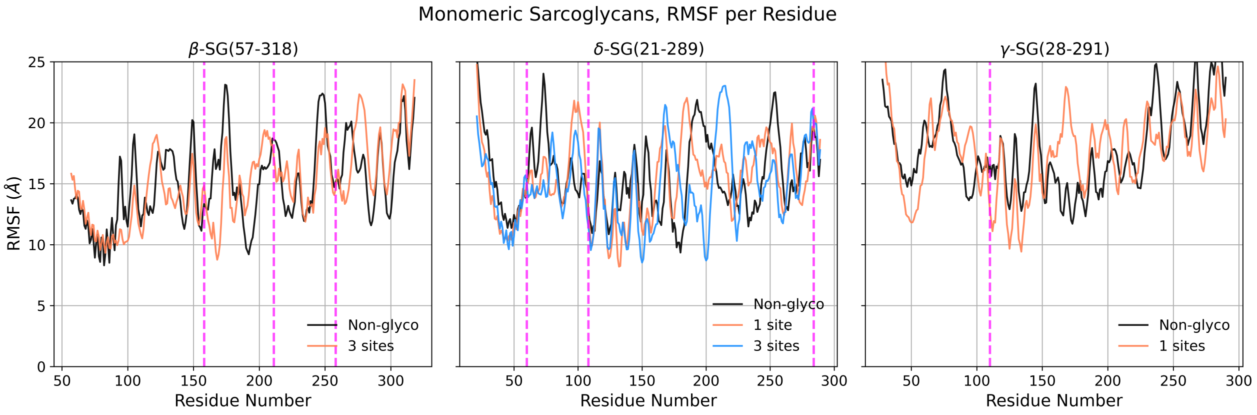


Figure S2: RMSF of SG subunits. In each subunits, $C_\alpha$ of residues was used to calculate the RMSF. The plot on the left belongs to $\beta$-SG, the one in the middle to $\delta$-SG and the one on the right to $\gamma$-SG.

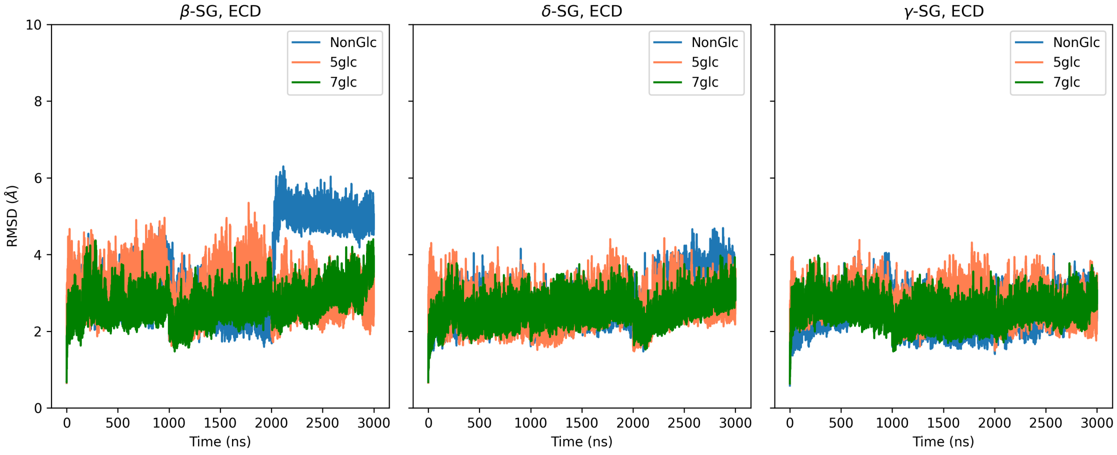


Figure S3: RMSD comparison between glycosylated (5- and 7-site: 5Glc and 7Glc, respectively) and nonglycosylated (NonGlc) forms of the SG complex over a 3$\mu$s accumulated trajectory. The right panel shows $\beta$-SG, the middle panel $\delta$-SG, and the left panel $\gamma$-SG components of the complex.

Table S4: RMSF differences at the glycosylation site in isolated SG subunits.
$\Delta RMSF_i = RMSF_{i,P} - RMSF_{i,GP}$ , where $i$ is for the glycosylated site, **GP** for glycosylated protein, and **P** for the nonglycosylated protein.

| Name | # of glycans | Glycosylation site | # of sugar residues | Location of glycosylation site | $\Delta RMSF_i$ | Avg. \| ΔRMSF \| |
|---|---|---|---|---|---|---|
| | | ASN-158 | 2 | loop | 1.4812 | |
| β-SG | 3 | ASN-211 | 3 | loop | -1.3958 | 1.1478 |
| | | ASN-258 | 3 | β-sheet | -0.5664 | |
| | | ASN-108 | | loop | -0.6662 | |
| δ-SG | 1 | - | 3 | loop | - | - |
| | | - | | loop | - | |
| | | ASN-60 | 3 | loop | -0.0446 | |
| δ-SG | 3 | ASN-108 | 3 | loop | -1.4335 | 0.5337 |
| | | ASN-284 | 3 | loop | 0.1231 | |
| γ-SG | 1 | ASN-110 | 3 | loop | -3.6695 | - |

Table S5: Glycosylation site information and RMSF differences at the glycosylation site, when 5 of 7 reported glycosylation sites are glycosylated.
$\Delta RMSF_i = RMSF_{i,P} - RMSF_{i,GP}$ , where $i$ is for the glycosylated site, **GP** for glycosylated protein, and **P** for the nonglycosylated protein.

| Name | # of glycans | Glycosylation site | # of sugar residues | Location of glycosylation site | $\Delta RMSF_i$ | Avg. \| ΔRMSF \| |
|---|---|---|---|---|---|---|
| | | ASN-158 | 2 | loop | 0.1647 | |
| β-SG | 3 | ASN-211 | 3 | loop | 0.439 | 0.2232 |
| | | ASN-258 | 3 | β-sheet | 0.0661 | |
| | | ASN-108 | 3 | loop | -0.0065 | |
| δ-SG | 1 | - | - | - | - | - |
| | | - | - | - | - | |
| γ-SG | 1 | ASN-110 | 3 | loop | -0.0352 | - |

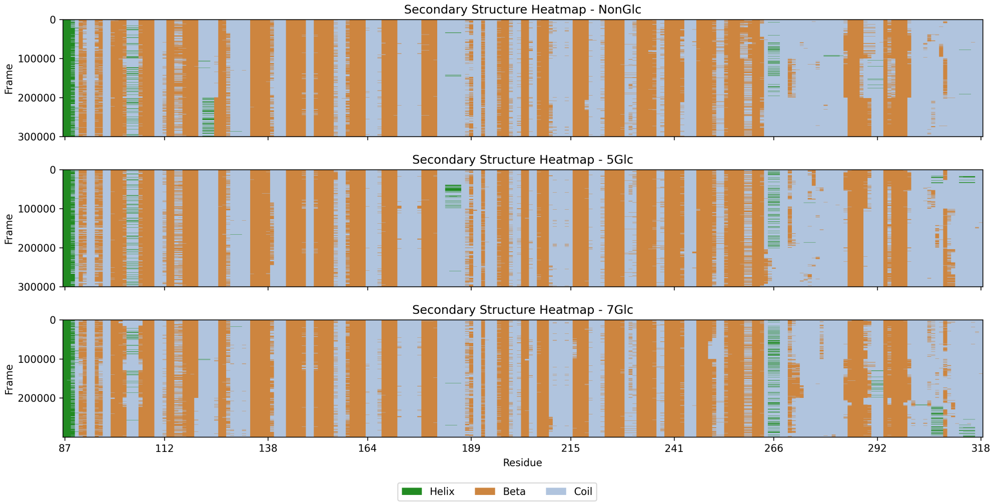


(a) ECD of $\beta$-SG in the SG complex

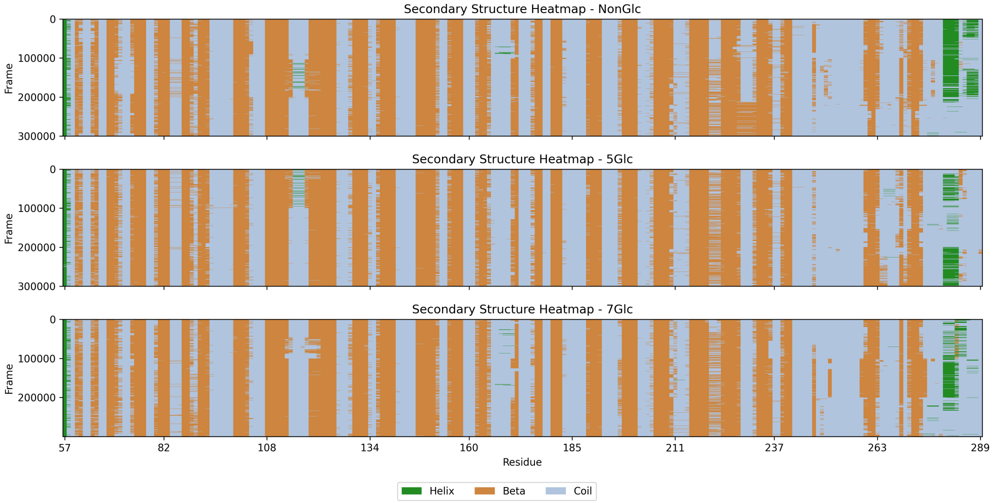


(b) ECD of $\delta$-SG in the SG complex

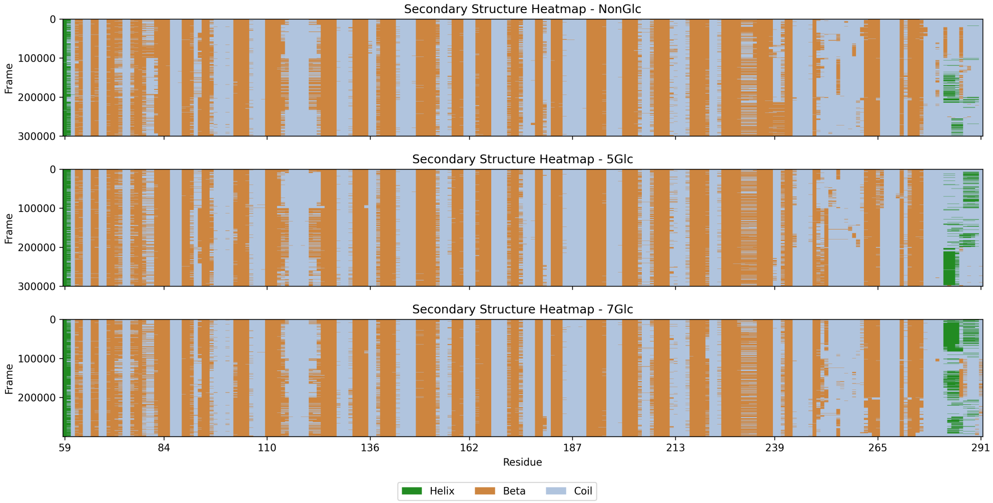


(c) ECD of $\gamma$-SG in the SG complex

Figure S4: Secondary structure (DSSP) analysis of the SG complex components under nonglycosylated and glycosylated (5Glc and 7Glc) conditions over 3$\mu$s of accumulated data.

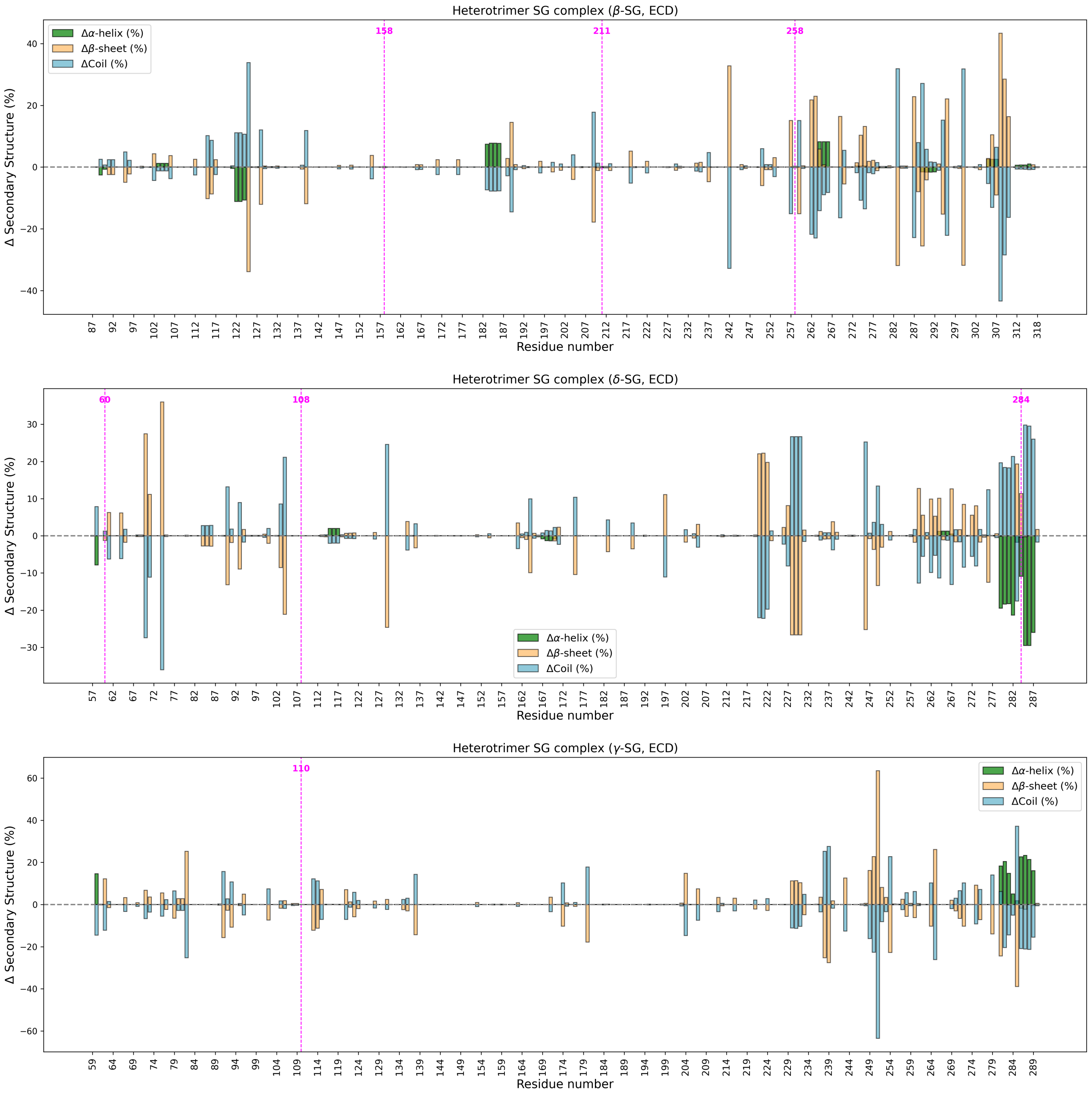


Figure S5: The differences in percentages between glycosylated (5 glycosylation sites in total) and non-glycosylated states were computed to quantify residue-level structural changes for components of SG complex ($\beta$-, $\delta$- and $\gamma$- SG). The magenta dashed lines show the glycosylation sites.

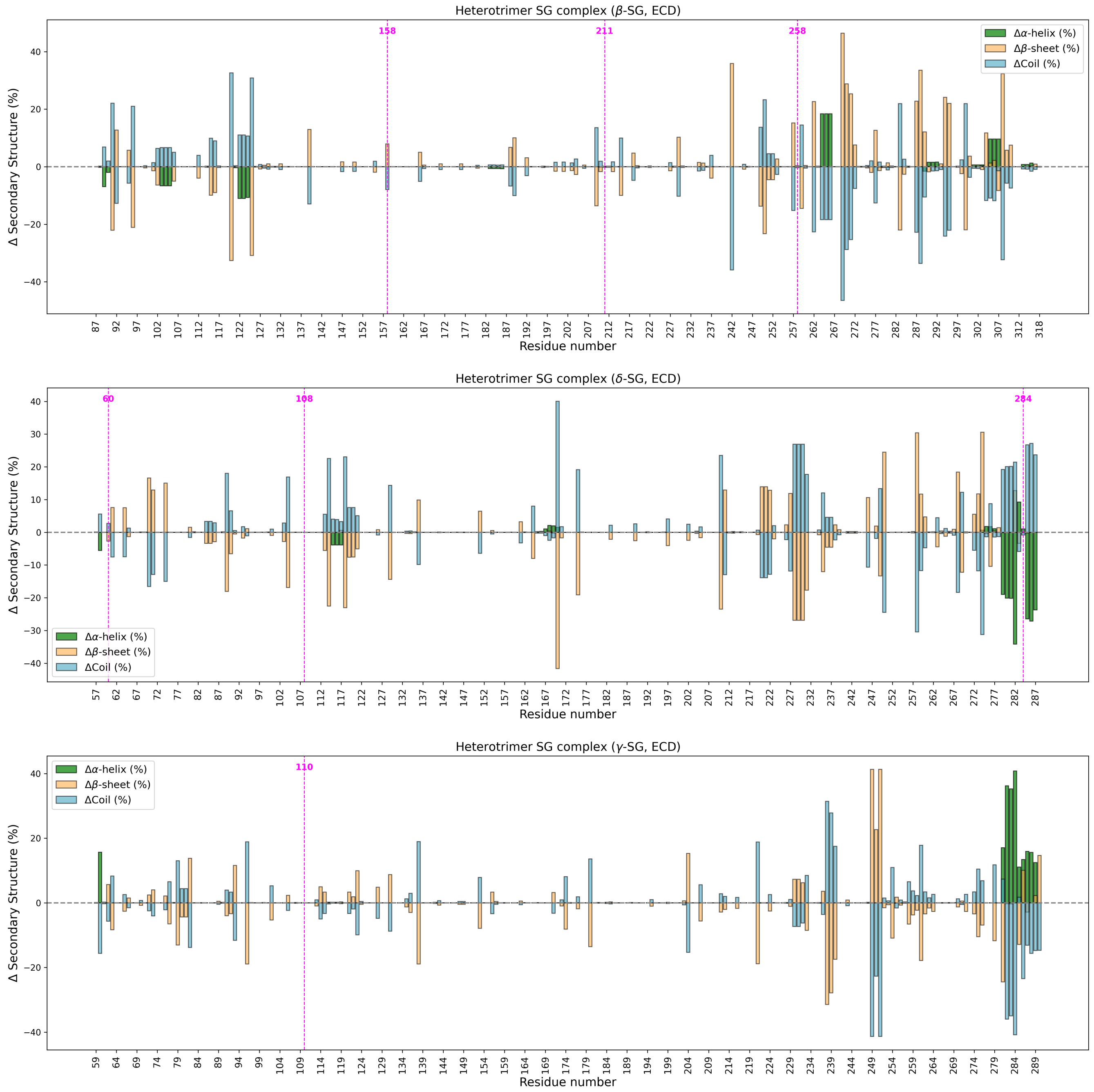


Figure S6: The differences in percentages between glycosylated (7 glycosylation sites in total) and non-glycosylated states were computed to quantify residue-level structural changes for components of SG complex ($\beta$-, $\delta$- and $\gamma$- SG). The magenta dashed lines show the glycosylation sites.

Table S6: Glycosylation site information and RMSF differences at the glycosylation site, when all 7 reported glycosylation sites are glycosylated.
$\Delta\text{RMSF}_i = \text{RMSF}_{i,P} - \text{RMSF}_{i,GP}$ , where $i$ is for the glycosylated site, **GP** for glycosylated protein, and **P** for the nonglycosylated protein.

| Name | # of glycans | Glycosylation site | # of sugar residues | Location of glycosylation site | $\Delta\text{RMSF}_i$ | Avg. $\mid \Delta\text{RMSF} \mid$ |
|---|---|---|---|---|---|---|
| $\beta$-SG | 3 | ASN-158 | 2 | loop | 0.1642 | 0.1299 |
| | | ASN-211 | 3 | loop | 0.2037 | |
| | | ASN-258 | 3 | $\beta$-sheet | 0.0218 | |
| $\delta$-SG | 3 | ASN-60 | 3 | loop | -0.1554 | 0.6590 |
| | | ASN-108 | 3 | loop | 0.0102 | |
| | | ASN-284 | 3 | loop | 1.8115 | |
| $\gamma$-SG | 1 | ASN-110 | 3 | loop | -0.1829 | - |
